# Single-molecule tracking in live *Yersinia enterocolitica* reveals distinct cytosolic complexes of injectisome subunits

**DOI:** 10.1101/211318

**Authors:** Julian Rocha, Charles Richardson, Mingxing Zhang, Caroline Darch, Eugene Cai, Andreas Diepold, Andreas Gahlmann

## Abstract

In bacterial type 3 secretion, substrate proteins are actively transported from the bacterial cytoplasm into the host cell cytoplasm by a large membrane-embedded machinery called the injectisome. Injectisomes transport secretion substrates in response to specific environmental signals, but the molecular details by which the cytosolic secretion substrates are selected and transported through the type 3 secretion pathway remain unclear. Secretion activity and substrate selectivity are thought to be controlled by a sorting platform consisting of the proteins SctK, SctQ, SctL, and SctN, which together localize to the cytoplasmic side of membrane-embedded injectisomes. However, recent work revealed that sorting platform proteins additionally exhibit substantial cytosolic populations and that SctQ reversibly binds to and dissociates from the cytoplasmic side of membrane-embedded injectisomes. Based on these observations, we hypothesized that dynamic molecular turnover at the injectisome and cytosolic assembly among sorting platform proteins is a critical regulatory component of type 3 secretion. To determine whether sorting platform complexes exist in the cytosol, we measured the diffusive properties of the two central sorting platform proteins, SctQ and SctL, using live cell high-throughput 3D single-molecule tracking microscopy. Single-molecule trajectories, measured in wild-type and mutant *Yersinia enterocolitica* cells, reveal that both SctQ and SctL exist in several distinct diffusive states in the cytosol, indicating that these proteins form stable homo- and hetero-oligomeric complexes in their native environment. Our findings provide the first diffusive state-resolved insights into the dynamic regulatory network that interfaces stationary membrane-embedded injectisomes with the soluble cytosolic components of the type 3 secretion system.

## Introduction

Bacteria can secrete intact macromolecules into the surrounding medium or into nearby cells through broadly-conserved large biomolecular assemblies, called secretion systems (1). Type 3 secretion systems (T3SSs), in particular, are used by bacteria for flagellar biogenesis and for virulence (2). The virulence-associated T3SS, also called the injectisome, features a long hollow needle that protrudes from the bacterial cell surface and ultimately anchors itself into the host cell membrane (**Fig. 1**) (3–7). T3SS injectisomes target host cell biology through the translocation of species-specific effector proteins into the host cell cytoplasm. The structural proteins of the T3SS injectisome are highly conserved among prominent Gram-negative pathogens, including *Salmonella, Shigella, Yersinia, Pseudomonas*, and *E. coli*, making T3SSs broadly utilized protein delivery machines.

**Fig. 1.**
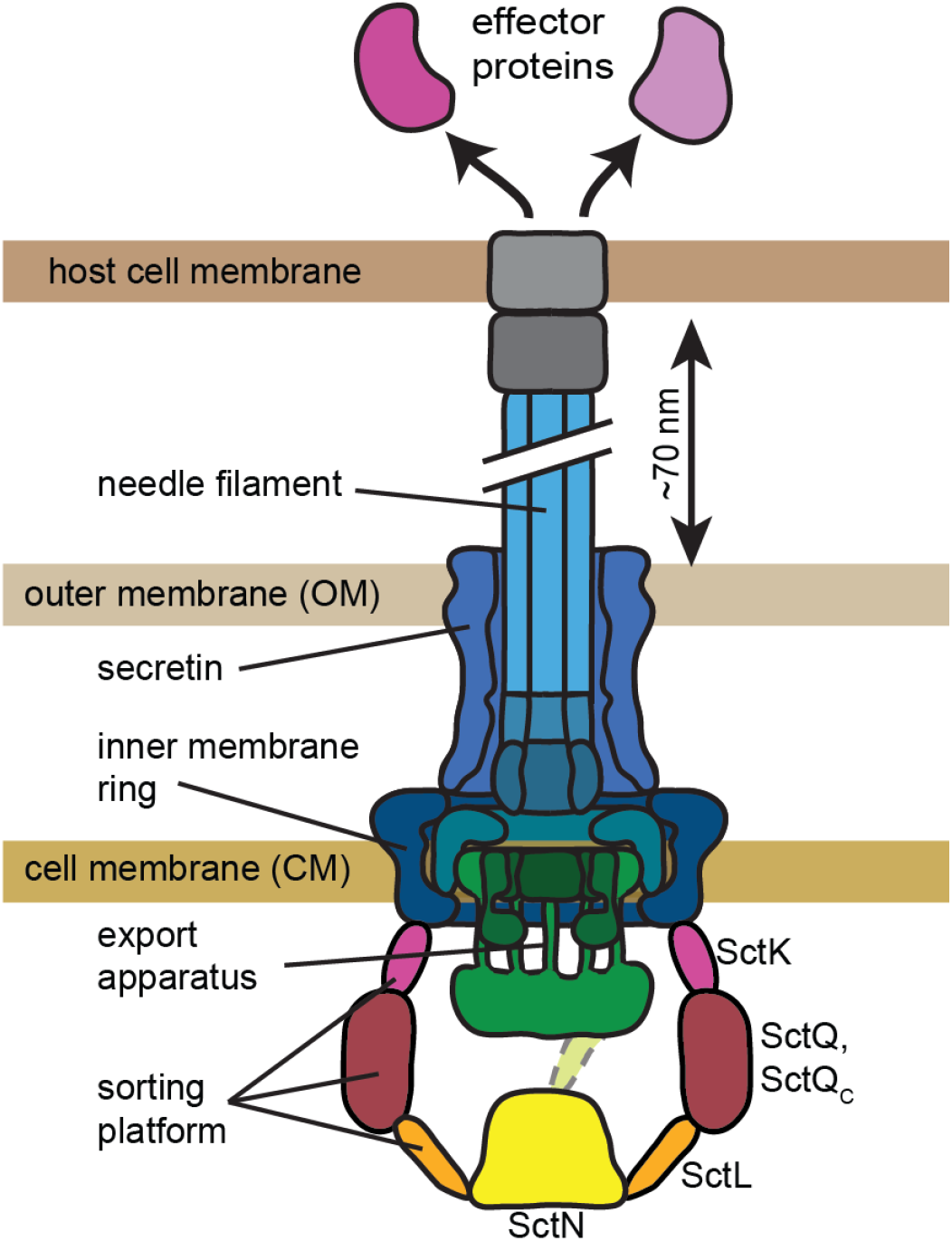
The T3SS injectisome spans both the inner and outer bacterial membranes and features a long hollow needle that protrudes away from the cell surface. The injectisome is used to transport effector proteins from the bacterial cytosol into the cytosol of eukaryotic host cells. At the cytoplasmic interface of the fully assembled injectisome, three interacting proteins (SctL,Q,K) are essential for the function of the T3SS and together form a so-called sorting platform. SctL forms a cradle-like structure that connects the hexameric ATPase SctN to each of the six pods, which contain multiple SctQ (and likely SctQC) subunits (21). SctK is an auxiliary protein that resides between the SctQ pods and the inner membrane ring of the injectisome. Figure adapted from Ref. (54).

A key feature of virulent T3SSs is that secretion can be activated in response to environmental cues and that substrate selectivity follows a well-defined temporal hierarchy (6). Expression and assembly of the injectisome is triggered by a temperature jump to 37°C experienced upon entry into mammalian hosts(3). Early substrates of the secretion-competent injectisome are proteins that assemble the needle itself. Once the needle has reached a certain length, selectivity is switched to middle secretion substrates that form the translocon pore, which anchors the needle tip into the host cell membrane. Finally, selectivity switches again upon stable injectisome-mediated host cell contact (8) or in response to external cues, such as calcium chelation in *Yersinia* (9), to allow secretion of late secretion substrates, namely the effector proteins (5).

A prominent model of T3SS functional regulation posits that selection of different export substrates is enabled through coordinated interactions among the cytoplasmic components of the injectisome (**Fig. 1**) (10, 11). Interactions between SctK, SctQ, SctL, and SctN (YscK, YscQ, YscL, and YscN in *Yersinia*; the universal nomenclature is used throughout this manuscript) are indeed essential for type 3 secretion (12–17) and for their mutual localization to the injectisome (18–20). Importantly, proteomic experiments isolated SctK, SctQ, SctL, and SctN as soluble high molecular weight complexes (16). Different secretion substrates and their cognate cytosolic chaperones were also found in high molecular weight complexes containing SctQ, SctK, and SctL (11, 16). These observations suggest that SctK:Q:L:N complexes function as cytoplasmic sorting platforms that control the order of protein secretion.

The precise structure of the sorting platform proteins SctK, SctQ, SctL, and the ATPase SctN within fully assembled injectisomes has recently been determined by cryo-electron tomography (10, 21, 22). The 3D tomogram averages reveal a cytoplasmic injectisome complex of hexametric symmetry, which is notably different from the continuous cytoplasmic ring (C-ring) structure observed in flagellar T3SSs (23–25). The cytoplasmic complex of the injectisome consists of six “pods” containing SctQ, presumably in complex with additional copies of its C-terminus, SctQC, which is translated from an internal translation initiation site (26, 27). These pods attach to the membrane-embedded ring of the needle complex, presumably through SctK (21). Each pod further connects to one of six spokes of a cradle-like structure formed by SctL that holds in place the central hexameric SctN (**Fig. 1**). The static *in situ* morphologies provided unprecedented insight into how SctK, SctQ, SctL, and SctN are arranged relative to each other when bound to the injectisome, but the obtained results did not provide clues into how their sorting functionality could be realized.

Live-cell compatible approaches revealed that the quaternary structure of the cytoplasmic injectisome components is highly dynamic. Fluorescence recovery after photobleaching (FRAP) measurements in *Y. enterocolitica* showed that the sorting platform protein SctQ continuously exchanges between an injectisome bound state and a freely diffusing cytosolic state and that the exchange rate doubled upon chemical activation of protein secretion (18). Other sorting platform proteins, SctK, SctL, and SctN, also exhibit substantial cytosolic populations and fluorescence correlation spectroscopy (FCS) measurements revealed that their diffusion coefficients were altered in different deletion mutants and between secreting and non-secreting conditions (19). These observations suggest that fully-assembled sorting platforms may be natively present in the cytosol as freely diffusing complexes that are functionally relevant for secretion. However, so far, it was not clear whether well-defined cytosolic complexes are responsible for the observed effects. Here we show formation of distinct homo- and hetero-oligomeric cytosolic complexes of sorting platform proteins using high-throughput 3D single-molecule tracking measurements in live *Y. enterocolitica.* Our results demonstrate that the two central sorting platform proteins, SctQ and SctL, interact with each other and with other T3SS proteins not just at the injectisome, but also in the cytosol, resulting in the formation of several distinct molecular complexes. We further show that the relative population fractions of these complexes is dependent on the presence of other T3SS proteins and changes with type 3 secretion activity. These results suggest that functional regulation of T3SS may occur away from the membrane-embedded injectisomes through the ordered formation of distinct complexes in the bacterial cytosol.

## Experimental Procedures

### Bacterial Strains and Plasmids

*Yersinia enterocolitica* strains were generated by allelic exchange as previously described (20, 28). Mutator plasmids harboring 250-500 bp flanking regions, the coding sequences of eYFP or PAmCherry1, and a glycine-rich 13 amino acid linker between the fluorescent protein and the target protein were introduced into *E. coli* SM10 λpir for conjugation with *Y. enterocolitica* pIML421asd (29). After sucrose counter-selection for the second allelic exchange event, fluorescent *Y. enterocolitica* were analyzed by PCR to confirm target insertion.

Plasmids for the inducible exogenous expression of fluorescent and fluorescently-tagged proteins were derived from IPTG-inducible pAH12 and arabinose-inducible pBAD vectors. The coding sequences of eYFP and PAmCherry1 were PCR amplified using Q5 DNA polymerase (New England Biolabs, Ipswich, Maine) from pXYFPN-2 and pVPAmCHY-popZ, respectively (30). The PCR product was isolated using a gel purification kit (Invitrogen, Carlsbad, California) and used as a megaprimer for amplification and introduction into a pAH12-derivative containing a kanamycin resistance cassette, LacI, and a lac promoter to generate pAH12-eYFP and pAH12-PAmCherry1. The pAH12 backbone was a gift from Carrie Wilmot. A series of pBAD expression vectors for eYFP, eYFP-SctQ, eYFP-SctQM218A, PAmCherry1-SctL, and PAmCherry1 were generated from mVenus-pBAD, originally developed by Michael Davidson (Addgene, Cambridge, Massachusetts, plasmid #54845). The coding sequence for eYFP was amplified from pAH12-eYFP. The coding sequence for eYFP-SctQ was amplified from *Y. enterocolitica* strain AD4442 [eYFP-SctQ]. The eYFP-SctQM218A coding sequence variant was generated using piecewise PCR of AD4442 with both the 5’ and 3’ fragments containing sequences overlapping the coding region corresponding to the M218A mutation. These fragments were gel purified and combined using outside primers with Q5 DNA polymerase. The coding sequence for PAmCherry1-SctL was amplified from *Y. enterocolitica* strain AD4459. The coding sequence for PAmCherry1 was amplified from pAH12-PAmCherry1. All final PCR products were created with Q5 DNA polymerase and gel purified. Purified products were incubated with Taq DNA polymerase (Thermo Scientific, Waltham, Massachusetts) and dNTPs at 72°C. PCR reactions were TA cloned using pCR2.1-TOPO (Invitrogen) according to the manufacturer’s directions. After screening for insert using Taq DNA polymerase, plasmids from positive clones were isolated using a miniprep kit (Omega Biotek, Norcross, Georgia). mVenus-pBAD and pCR2.1 minipreps were digested with EcoRI and XhoI restriction enzymes (New England Biolabs). Digested vector and inserts were ligated using T4 DNA ligase and transformed into *E. coli* TOP10 cells. Colonies were PCR screened for presence of correct insert. All plasmids were sequenced by GeneWiz (South Plainfield, New Jersey) prior to electroporation into *Y. enterocolitica* for analysis. A list of all strains and plasmids can be found in **Table S1**.

### Cell Culture

*Y. enterocolitica* cultures were inoculated from a freezer stock in BHI media (Sigma Aldrich, St. Louis, Missouri) with nalidixic acid (Sigma Aldrich) [35 μg/mL] and 2,6-diaminopimelic acid (Chem Impex International, Wood Dale, Illinois) [80 μg/mL] one day prior to an experiment and grown at 28°C with shaking. After 24 hours, 300 μL of overnight culture was diluted in 5 mL fresh BHI, nalidixic acid, and diaminopimelic acid (dap) and grown at 28°C for another 60-90 minutes. For imaging cells in the secretion ON state, glycerol [4 mg/mL], MgCl2 [20 mM] and EDTA [5 mM] were added to the culture medium. For imaging cells in the secretion OFF state, glycerol, MgCl_2_, and CaCl2 [5 mM] were added to the culture medium. In both cases, the *yop* regulon was induced by rapidly shifting the cultures to 37°C in a water bath(9), and cells were incubated at 37°C with shaking for another 3 hours prior to imaging. After induction, cells were harvested by centrifugation at 5000 g for 3 minutes and washed 3 times with M2G (4.9 mM Na2HPO4, 3.1 mM KH2PO4, 7.5 mM NH_4_Cl, 0.5 mM MgSO4, 10 μM FeSO4 (EDTA chelate; Sigma), 0.5 mM CaCl_2_) with 0.2% glucose as the sole carbon source). The remaining pellet was then re-suspended in M2G, dap, MgCl_2_, glycerol, and EDTA/CaCl_2_. Cells were plated on 1.5 – 2% agarose pads in M2G containing dap, glycerol, and MgCl_2_.

Plasmids were introduced into *Y. enterocolitica* cells using electroporation. Transformed cells were plated on LB agar [10 g/L peptone, 5 g/L yeast extract, 10 g/L NaCl, 1.5% agar] (Fisher Scientific, Hampton, New Hampshire) containing kanamycin [50 μg/mL] or ampicillin [100 μg/mL] for cells containing pAH12-or pBAD-derived plasmids, respectively. For electroporation of *Y. enterocolitica* pIML421asd cells, recovery media and plates also contained dap. Plasmid containing cells were inoculated similarly, except inoculation media also contained kanamycin or ampicillin for pAH12- or pBAD-based plasmids, respectively. Prior to imaging cell cultures were rapidly temperature shifted to 37°C and incubated for 3 hours. Cultures of cells containing pAH12- or pBAD-based plasmids were induced with IPTG (Sigma Aldrich) [0.2 mM, final] or arabinose (Chem Impex) [0.2%], respectively, for the final 2 hours of incubation.

### Secretion Assay and Immunoblot

Cultures for protein secretion assays and immunoblot analysis were inoculated to an optical density at 600 nm (OD600) of 0.15 in BHI supplemented with 35 μg/ml nalidixic acid, 80 μg/ml diaminopimelic acid, 0.4% glycerol, 20 mM MgCl_2_, and 5 mM EDTA. Cultures were agitated at 28°C for 90 min. The *yop* regulon was then induced by shifting the temperature to 37°C in a water bath, where cultures were agitated for another 180 min. Supernatant and whole cells were separated by centrifugation (5 min, 21,000 g). Secreted proteins were precipitated with 10% trichloroacetic acid overnight at 4°C. An equivalent of proteins secreted by 3·10^8^ bacteria was used for further analysis of the supernatant, whereas the lysate of the equivalent of 10^8^ bacteria was loaded onto the gel for analysis of total cellular proteins. Proteins were separated on Novex 4-20% gradient SDS–PAGE gels and stained using the Coomassie-based ‘Instant blue’ staining solution (Expedeon, San Diego, California), or immunoblotted using rabbit polyclonal antibodies against *Y. enterocolitica* SctQ (MIPA235; 1:1,000) (**Fig. S1**).

### Super-resolution Fluorescence Imaging

Experiments were performed on a custom-built dual-color inverted fluorescence microscope based on the RM21 platform (Mad City Labs, Inc, Madison, Wisconsin). Immersion oil was placed between the objective lens (UPLSAPO 100X 1.4 NA) and the glass cover slip (VWR, Radnor, Pennsylvania, #1.5, 22mmx22mm). Single-molecule images were obtained by utilizing eYFP photoblinking (31) and PAmCherry1 photo-activation (32). A 514 nm laser (Coherent, Santa Clara, California, Genesis MX514 MTM) was used for excitation of eYFP (~350 W/cm^2^) and 561nm laser (Coherent Genesis MX561 MTM) was used for excitation of PAmCherry1 (~350 W/cm^2^). A 405 nm laser (Coherent OBIS 405nm LX) was used to photo-activate PAmCherry1 (~20 W/cm^2^) simultaneously with 561nm excitation. Zero order quarter wave plates (Thorlabs, Newton, New Jersey, WPQ05M-405, WPQ05M-514, WPQ05M-561) were used to circularly polarize all excitation lasers, and the spectral profile of the 514nm laser was filtered using a bandpass filter (Chroma, Bellows Falls, Vermont, ET510/10bp). Fluorescence emission from both eYFP and PAmCherry1 was passed through a shared filter set (Semrock, Rochester, New York, LP02-514RU-25, Semrock NF03-561E-25, and Chroma ET700SP-2P8). A dichroic beam splitter (Chroma T560lpxr-uf3) was then used to split the emission pathway into ‘red’ and ‘green’ channels. An additional 561nm notch filter (Chroma ZET561NF) was inserted into the ‘red’ channel to block scattered laser light. Each emission path contains a wavelength specific dielectric phase mask (Double Helix, LLC, Boulder, Colorado) that is placed in the Fourier plane of the microscope to generate a double-helix point-spread-function (DHPSF) (33, 34). The fluorescence signals in both channels are detected on two separate sCMOS cameras (Hamamatsu, Bridgewater, New Jersey, ORCA-Flash 4.0 V2). Up to 20,000 frames are collected per field-of-view with an exposure time of 25ms. A flip-mirror in the emission pathway enables toggling the microscope between fluorescence imaging and phase contrast imaging modes without having to change the objective lens of the microscope.

### Data Processing

Raw image processing and analysis was carried out using MATLAB (The MathWorks, Inc, Natick, Massachusetts) with a modified version of the easyDHPSF code (35). Maximum likelihood estimation using a double-Gaussian PSF model was used to extract the 3D localizations of single-molecule emitters from raw images collected on sCMOS cameras (36). For background subtraction, a median filter with a time window of 10 frames was used (37).

Single molecule localizations were assigned to individual cells based on the corresponding phase contrast image. Cell outlines were generated based on the phase contrast images using the open-source software OUFTI (38). The outlines are registered to the fluorescence data by a two-step 2D affine transformation using the ‘cp2tform’ function in MATLAB. In the first step, five control point pairs were manually selected by estimating the position of the cell poles based on single-molecule localization data and the cell outlines generated by OUFTI. An initial transformation was generated, and cell outlines containing less than 10 localizations were removed. The center of mass for all remaining cell outlines and single-molecule localizations within them were then used to generate a second, larger set of control point pairs to compute the final transformation function. A large set of control points (N ~ 100 cells) ensures that cells with few localizations or cells positioned partly outside the field-of-view do not skew the transformation. Only localizations that lie within the cell outlines were considered for further analysis. Analysis of high density regions of localizations was carried out using the density based clustering algorithm DBSCAN (39), using a distance parameter of 75 nm and a minimum number of points parameter of 15.

### Single Molecule Tracking Analysis

To determine the apparent diffusion coefficients of single molecules, 3D single-molecule localizations in subsequent frames were linked into trajectories using a distance threshold of 2.5 μm. Only trajectories with at least 4 localizations were considered for further analysis. In addition, if two or more localizations were present in the cell at the same time during the length of the trajectory, the trajectory was not considered for further analysis. These steps minimized the linking problem, in which, due to misassignment, two or more molecules could contribute to the same trajectory (40).

The Mean Squared Displacement (MSD) was calculated according to

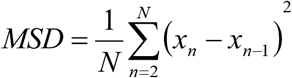

where *N* is the total number of time points and *x_n_* is the position of the molecule at time point *n*. The apparent diffusion coefficient, *D*\*, of a given single-molecule was then computed as

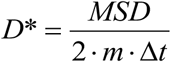

where *m* is the dimensionality of the problem and *Δt* is the camera exposure time. In our experiments m=3 and *Δt* =25 ms. It is important to note that the so-estimated single-step apparent diffusion coefficients do not take into account the static and dynamic localization errors (41), or the effect of confinement within the bacterial cells. Instead of accounting for these effects using analytical models (42, 43), we generated simulated noise and motion-blurred images of diffusing molecules in rod-shaped cell volumes, as described in the following section. The resulting images were then analyzed in the same manner as experimental data. These steps ensured that static and dynamic localization errors and the effect of confinement within the bacterial cells are accurately modeled.

### Monte Carlo Simulations

To resolve the unconfined diffusion coefficients of predominant molecular complexes in living cells based on the experimentally measured distribution of apparent diffusion coefficients, we performed Monte Carlo simulations of confined Brownian motion inside the volume of a cylinder using a set of 64 diffusion coefficients ranging from 0.05–20 μm^2^/s as input parameters. The size of the confining cylinder was chosen to match the average size of a *Y. enterocolitica* cell (radius = 0.4 μm, length = 5 μm). The starting position of the trajectory was randomly set within the volume of the cylinder and Brownian motion was simulated using short time intervals of 100 ns. We assumed a hard sphere reflection at the cell boundaries, i.e. if a molecule was displaced outside of the volume of the cylinder within a time step, it was redirected back towards the inside of the cylinder at a random angle. Choosing a short time step further ensured that the entire volume of the cylinder could be sampled by the diffuser, even the interfacial region near the cell boundary.

To simulate raw experimental data, we generated noisy, motion-blurred single-molecule images. Specifically, we generated DHPSF images corresponding to 50 periodically sampled positions of a molecule during the camera exposure time (25ms) and then summed these 50 sub images to obtain the motion-blurred DHPSF images, which when analyzed will produce position estimates with limited accuracy and precision due to dynamic localization error (41). The photon count of each simulated image was scaled to match the experimentally measured distribution for eYFP photon counts and a laser background of ~13 photons/pixel was added. Poisson noise was added to the image based on final photon count in each pixel. Finally, a dark offset (50 photons/pixel on average) with Gaussian read noise (σ ~1.5 photons) was added to produce the final image. This image was then multiplied by the experimentally measured pixel-dependent gain of our sCMOS camera to obtain an image in units of detector counts. These simulated images were then processed the same way as experimental data to obtain single-molecule localization, which were then linked into trajectories. Simulated trajectories were limited to six displacement steps to match the average length of our experimentally measure trajectories.

We simulated *N* = 5000 trajectories for each of the 64 input diffusion coefficients to obtain an array of *N* apparent diffusion coefficients (**Fig. S2 C and D**). The empirical cumulative distribution functions (eCDFs) corresponding to the 64 diffusion coefficients were then interpolated using a B-spline (order 25) and normalized individually. The interpolated array of 64 CDFs was then interpolated again along the *D* dimension using the ‘scatteredInterpolant’ function in MATLAB. This two-step interpolation provides a continuous 2-dimensional function that can then be used to compute the apparent diffusion coefficient distribution that we would observe in our chosen confinement geometry for any true diffusion coefficient value in the range of 0.05 and 20 μm^2^/s. This approach revealed that the experimentally measured apparent diffusion coefficients in *Y. enterocolitica*, are systematically decreased by up to 60% compared to unconfined diffusion (**Fig. S2A**). The simulated CDFs account for the effects of molecular confinement due to the cell boundaries, signal integration over the camera exposure time, as well as experimentally calibrated signal-to-noise levels.

### Data Fitting

Experimental distributions of apparent diffusion coefficients were fit using a linear combination of simulated CDFs, where each CDF corresponds to a single diffusive state that is described by a single diffusion coefficient (**Fig. S3**). Using the CDF for fitting instead of a histogram that represent the probability density function (PDF) eliminates bin-size ambiguities that can bias the fitting results. This approach allowed us to estimate the population fraction and the unconfined diffusion coefficient of each diffusive state that would have been obtained in the absence of cell boundaries. To obtain an initial estimate of the contributing diffusive states and their population fractions, constrained linear least-squares fitting was performed using the ‘lsqlin’ function in MATLAB and a periodically sampled array of simulated CDFs. This step allowed us to fit the leftmost portion of the experimentally measured empirical CDF (corresponding to stationary trajectories) using a sum of states with *D* < 0.5 μm^2^/s, which was then fixed for further analysis.

The distribution of estimated population fractions obtained from linear least squares fitting was assessed by bootstrapping (*N* = 100 samples) and any diffusive states that were not clearly resolved were combined into a single diffusive state. This approach resulted in initial values of diffusion coefficients of prevalent diffusive states and their respective population fraction) that were then refined further through non-linear least-squares fitting using the ‘fmincon’ function in MATLAB to obtain the optimal fitting parameter values reported in the main text. Confidence intervals for all fitting parameters were obtained by bootstrapping and are reported in **Table S2**.

To limit model complexity, the population fractions of diffusive states below 0.5 μm^2^/s were not refined through non-linear least-squares fitting, but instead assigned to stationary molecules. This choice was made because even completely stationary molecules exhibit non-zero apparent diffusion coefficients in single-molecule tracking experiments due to the non-zero single-molecule localization precision (static localization error). In our experiments, the *x*, *y*, and *z* localization precisions were 10-46 nm, 10-49 nm, and 16-71 nm, respectively (44). We confirmed that simulated trajectories of completely stationary emitters with these localization precisions produce apparent diffusion coefficients in the range of 0.01-0.15 μm^2^/s, which is in good agreement with the apparent diffusion coefficients estimated for clustered eYFP-SctQ localizations (**Fig. S5**).

### Radial Distribution Plotting

The radial distributions in **Fig. S6 and S7** were created using a combination of the cell outlines found with OUFTI and the localizations themselves. The localizations were separated using the cell outlines from OUFTI. The central axis of the cell was then found by projecting the localizations from sections of the cell along the cell long axis onto a 2D plane and finding the centroid of the localizations for each section. The radial distances were then found by calculating the distance of each 3D localization to the central axis.

## Results/Discussion

### Stationary SctQ localizes near the cell membrane

We introduced the coding sequence of eYFP in-frame near the translation start site of the SctQ coding sequence on the pYV virulence plasmid, which encodes all T3SS proteins in *Y. enterocolitica*, using previously described allelic replacement techniques (20, 28). In this way, the eYFP-SctQ fusion protein is expressed under the control of its native promoter. The cellular levels of eYFP-SctQ were increased compared to the native, unlabeled protein; we confirmed that the resulting fusion proteins did not result in detectable degradation products and were fully functional in an effector protein secretion assay (**Fig. S1**). We then used 3D-single molecule localization microscopy (3D-SMLM) to determine the subcellular positions of eYFP-SctQ molecules in live *Y. enterocolitica* under secreting conditions. The observed single-molecule localizations revealed clustering near the cell surface, and, when rendered as an intensity image, these clusters gave rise to the bright fluorescent foci (**Fig. 2 A and B**), which are similar to those observed by diffraction-limited and super-resolution microscopy(18, 20, 45). We used DBSCAN (39), a density-based clustering algorithm, to determine the positions and sizes of each cluster and found that clusters are preferentially localized within a 400-450 nm radius from the central cell axis (**Fig. 2 C and D**). This subcellular preference is consistent with membrane association, given the 469 ± 91 nm (mean ± s.d.) radius of *Y. enterocolitica* cells used in our experiments, and clusters have been previously observed to co-localize with other membrane-embedded injectisome components in two-color fluorescence imaging experiments(18, 20, 45). In contrast, cluster formation was not observed when we exogenously expressed eYFP-SctQ in a *Y. enterocolitica* strain lacking the pYV plasmid (pYV^−^) (**Fig. 2E**). Instead, eYFP-SctQ was uniformly distributed throughout the bacterial cytosol.

**Fig. 2.**
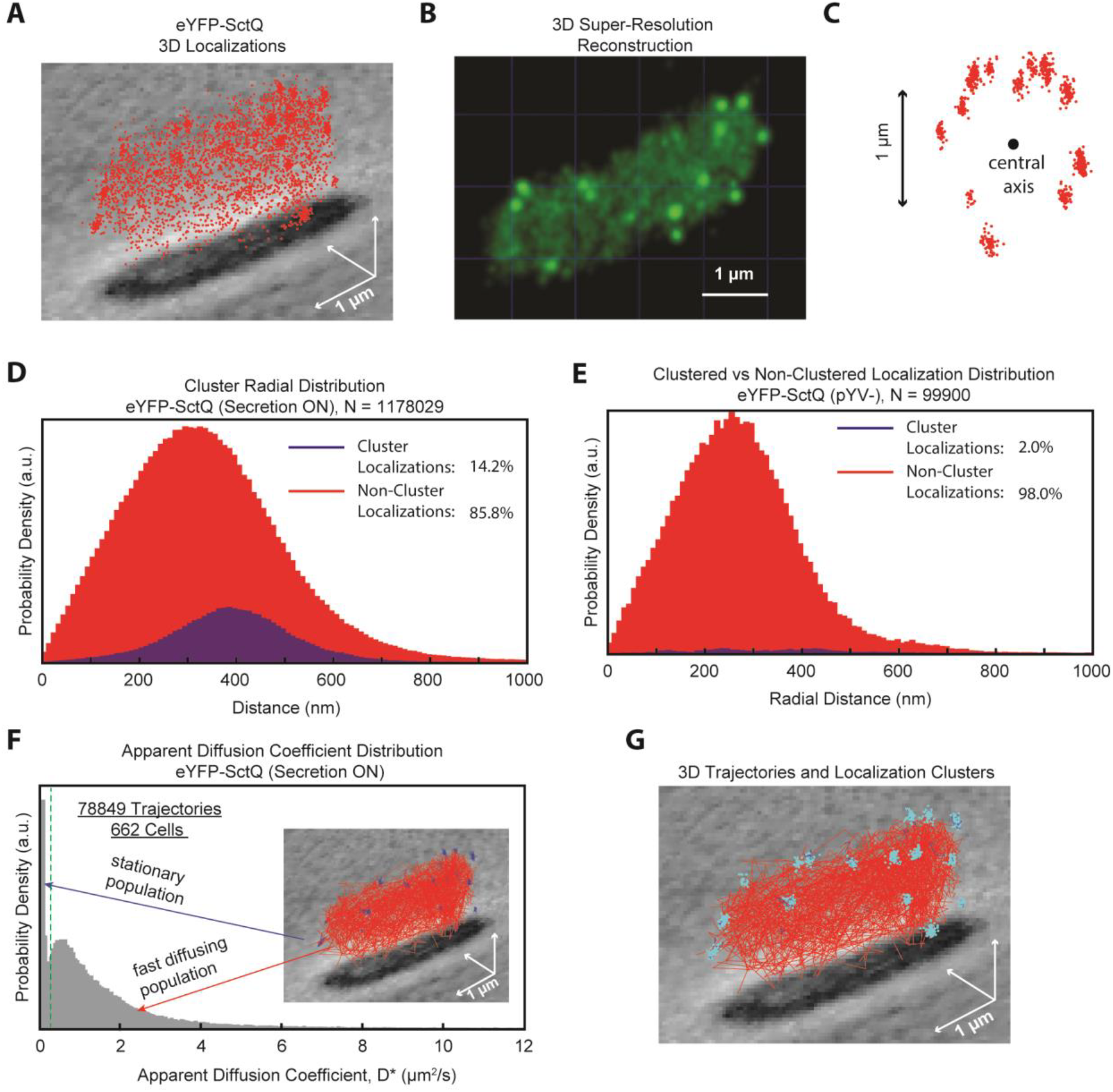
Fluorescently labelled eYFP-SctQ localizes throughout the cytosol and forms distinct clustered foci near the membrane. (A) Scatter plot of 3D localizations of eYFP-SctQ in a representative wild-type *Y. enterocolitica* cell overlaid on a phase contrast image of the cell. (B) Super-resolution image of the data shown in (A). Each single-molecule localization was rendered as a Gaussian sphere of width σ equal to the average localization precision. Closely overlapping spheres generate high intensity in the reconstructed image. An example cell for each data set can be found in **Fig. S8**. (C) Clustered localizations of eYFP-SctQ in a single cell, viewed along the central axis of the cell. (D) Radial distribution of clusters as determined by the DBSCAN clustering algorithm (39). Clustered localizations are enriched near the cell membrane. The distributions are broadened by cell width heterogeneity in the bacterial population and uncertainty in defining the central axis of the cells (E) 3D radial distribution of clusters for eYFP-SctQ in the pYV^−^ strain. Fewer clusters are present and do not preferentially localize near the cell membrane. (F) Probability density function of apparent diffusion coefficients for eYFP-SctQ. Two visually distinguishable populations emerge: a stationary (*D*\* < 0.15 μm^2^/s) and a mobile population (*D\** > 0.15 μm^2^/s). (F inset) Stationary (blue) and diffusive (red) trajectories in a single *Yersinia* cell. (G) Clustered localizations (teal) overlaid on the trajectory plot. Across the population (N = 662 cells) 60% of all stationary trajectories co-localized with clustered regions (within 100 nm of the cluster center-of-mass). The remaining 40% do not co-localize with clusters, because clustering algorithms rely user-defined parameters and are not 100% efficient in identifying clusters, especially in the presence of numerous diffuse localization, as is the case here.

Instead of only considering the localized molecules within clustered regions, as done by Zhang *et al*.,(45) we additionally quantified the diffusive properties of all localized molecules. Single-molecule tracking measurements in wild-type cells showed that the eYFP-SctQ population partitioned into a stationary and a mobile fraction (**Fig. 2F**). As expected, trajectories with slow apparent diffusion coefficients (*D*\* < 0.15 μm^2^/s) spatially co-localize with clusters (**Fig. 2G and Fig. S7A**), while the faster diffusing molecules (*D*\* > 0.15 μm^2^/s) localize randomly throughout the cell volume (**Fig. 2F inset, Fig. S7A**). The *D*\* = 0.15 μm^2^/s threshold was chosen based on the non-zero diffusion coefficients obtained for stationary emitters that are repeatedly localized with limited spatial localization precision (**Fig. S5**). We conclude that stationary membrane-embedded injectisomes serve as binding sites for SctQ molecules, and we assign the observed clusters or foci to injectisome-bound eYFP-SctQ molecules.

### SctQ exists in at least 3 diffusive states in the bacterial cytosol

The majority (~86%) of eYFP-SctQ localizations are not cluster-associated, but instead diffuse randomly throughout the cytoplasm (**Fig. 2 A, B, and D**). To examine the diffusive behaviors of unbound eYFP-SctQ, we determined the 3D trajectories of ~100,000 individual eYFP-SctQ molecules in hundreds of cells and computed the apparent diffusion coefficients for each trajectory (**Experimental Procedures**). As mentioned in the previous section, the resulting distribution of apparent diffusion coefficients shows two prominent peaks: one near ~0 μm^2^/s and the other at ~0.5 μm^2^/s (**Fig. 2F**). In addition to the peak at ~0.5 μm^2^/s, the distribution also shows a slow decay towards higher apparent diffusion coefficients. Using Monte Carlo simulations of anisotropically confined Brownian diffusion within rod-shaped bacterial cell volumes, we determined that such a distribution shape is only possible when multiple diffusive states manifest in the cell (**Experimental Procedures, Fig. S2**).

To estimate the unconfined diffusion coefficients (*D*) and population fractions of the diffusive states that are present in the cell, we used linear combinations of the Monte Carlo simulated apparent diffusion coefficient distributions to fit the experimentally measured distributions (Experimental Procedures). We found that fitting the eYFP-SctQ data required three cytosolic diffusive states corresponding to unconfined diffusion coefficients *D* = 1.1, 4.0, and 13.9 μm^2^/s with population fractions of 17%, 36%, and 22%, respectively (**Fig. 3 A and B, Table 1**). These components are in addition to a stationary component (*D* < 0.5 μm^2^/s) with population fraction of 24%. Confidence intervals for the optimized fitting parameters were obtained by bootstrapping and are reported in **Table S2**.

**Fig. 3.**
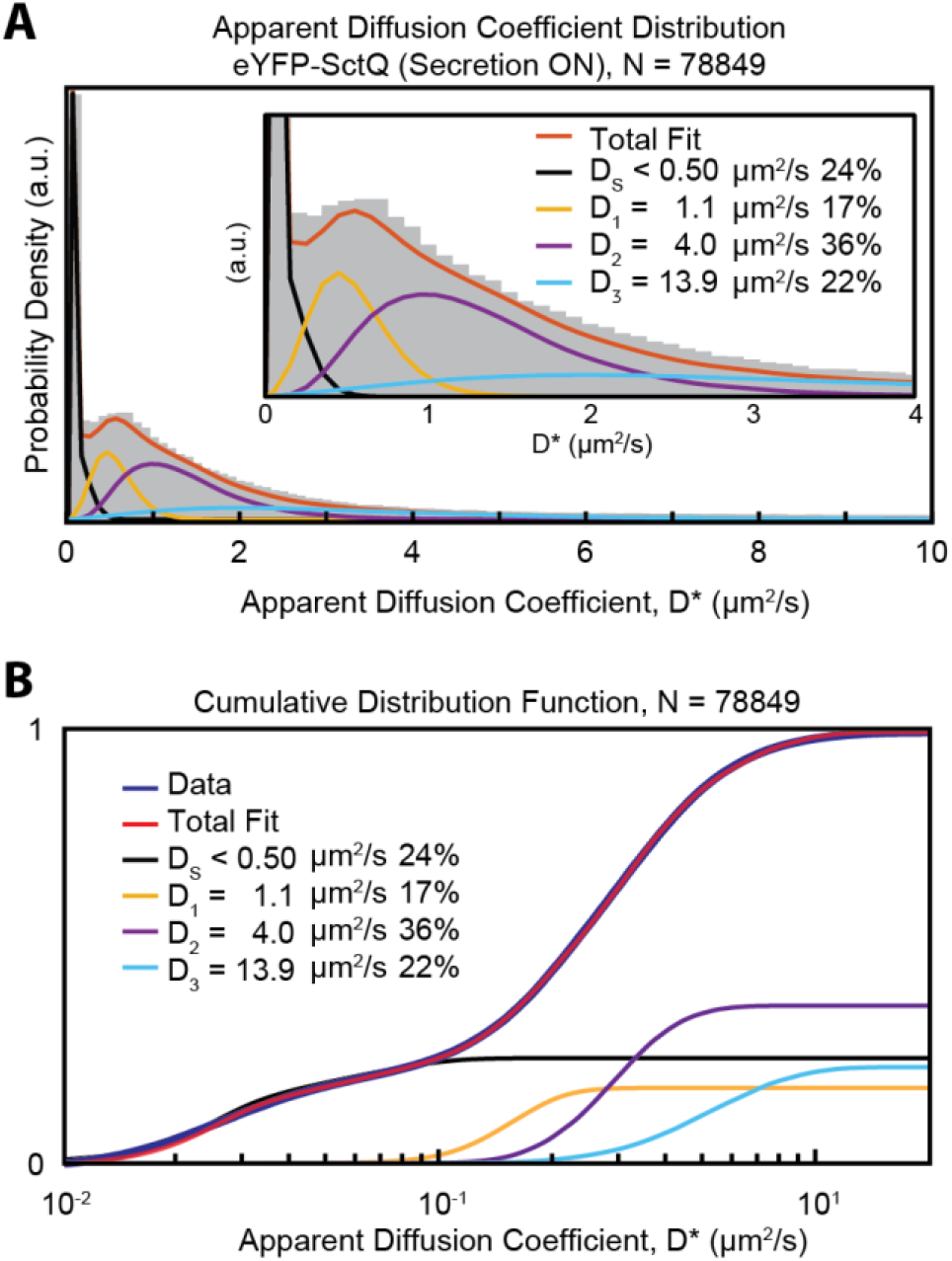
Fit of the distribution of apparent diffusion coefficients for eYFP-SctQ. (A) Fitting the distribution of eYFP-SctQ apparent diffusion coefficients required at least three diffusive states (D1-3) in addition to a stationary population (Ds). (B) The cumulative distribution function (CDF) of eYFP-SctQ apparent diffusion coefficients is fit using the indicated CDFs obtained by Monte-Carlo simulation of confined Brownian motion characterized by the indicated diffusion coefficients.

**Table 1.**
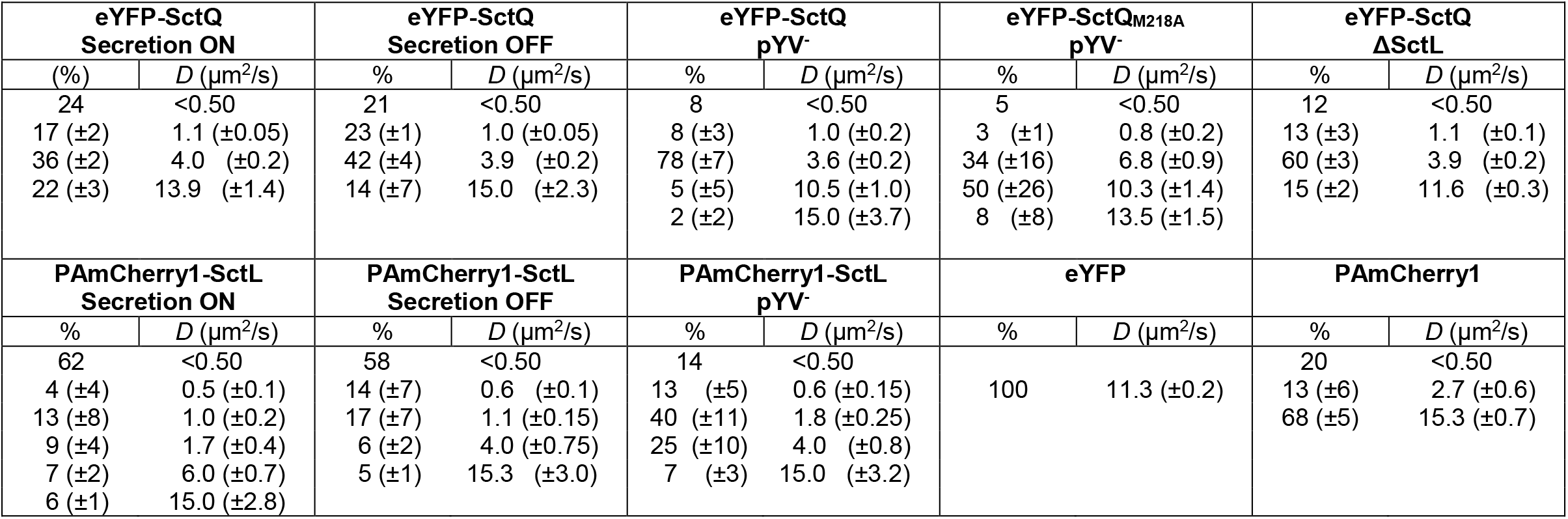
Fitted diffusion coefficients and relative population fractions.

As we did not detect any eYFP cleavage products (**Fig. S1**), the eYFP-SctQ monomer is the smallest and likely the fastest diffusing molecular species in our cells. Control experiments to measure the unconfined diffusion coefficients of eYFP confirmed that, when expressed in wild-type *Y. enterocolitica*, eYFP diffused at a similarly fast rate of *D* = 11.3 μm^2^/s (**Fig. S4I, Table 1**). We also observe a small stationary component for this protein suggesting that, even in the absence of known binding partners, the presence of immobile proteins cannot be ruled out in living cells. Assigning the fastest diffusive eYFP-SctQ state (D = 13.9 μm^2^/s) to the monomeric eYFP-SctQ fusion protein implies that the slower diffusive states at *D* ~ 1 and 4 μm^2^/s correspond to two distinct high molecular weight complexes, which could be eYFP-SctQ homo- or hetero-oligomers that involve additional T3SS proteins. To determine the molecular composition of these high molecular weight complexes, we performed further single-molecule tracking measurements in different genetic backgrounds.

### SctQ is capable of forming higher-order oligomers in the absence of other T3SS proteins

To test whether SctQ can form high molecular weight complexes in living cells independent of other T3SS proteins, we tracked single eYFP-SctQ molecules in a strain lacking all T3SS components (the pYV^−^ strain), for which membrane association was not observed (**Fig. 2E**). Fitting the distribution of apparent diffusion coefficients of eYFP-SctQ in the pYV^−^ strain required four diffusive states (**Fig. 4A, Table 1**). The predominant diffusive state (D = 3.6 μm^2^/s, 78%) is similar to the *D* = 4.0 μm^2^/s state observed for eYFP-SctQ in wild-type cells. These data show that formation of a specific oligomeric SctQ species is favored in pYV^−^ cells and does not require any other T3SS proteins. Conversely, the oligomerization behavior of eYFP-SctQ changes when other T3SS proteins are present, as there is a higher relative abundance of the putative eYFP-SctQ monomer in wild-type cells (22%) compared to pYV^−^ cells (~5%). If oligomerization of SctQ was completely unregulated we would expect a higher fraction of the *D* ~ 4 μm^2^/s diffusive state in wild-type cells, especially since the eYFP-SctQ fusion is express at slightly higher levels compared to the native, unlabeled protein (**Fig. S1**).

**Fig. 4.**
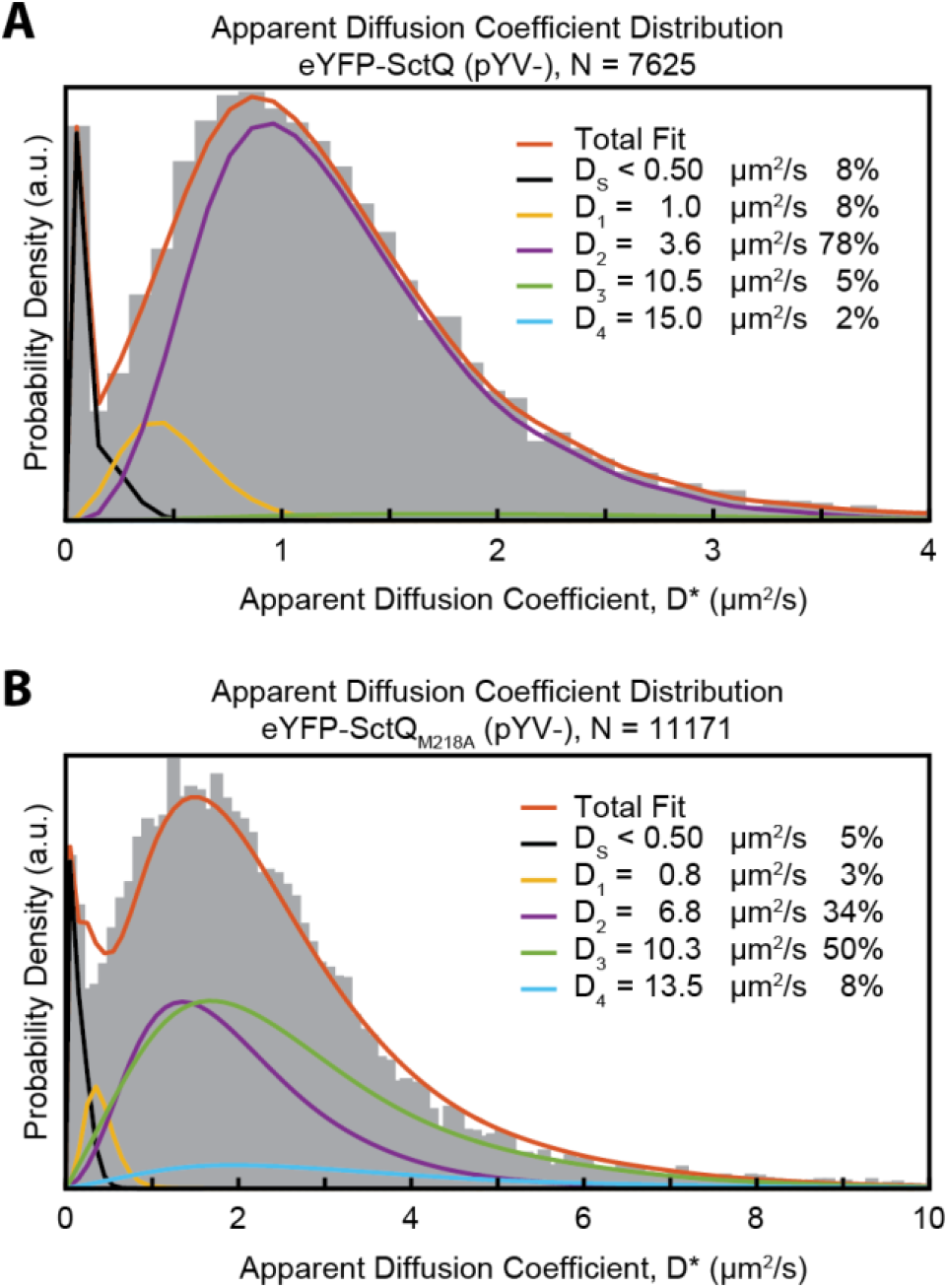
Fit of the apparent diffusion coefficient distributions for eYFP-SctQ in the absence of other T3SS proteins. (A) Fitting of the eYFP-SctQ apparent diffusion coefficients distribution in the pYV^−^ strain shows a dominant population at ~3.6 μm^2^/s. (B) A mutation in the SctQ coding sequence that suppresses the expression of SctQc eliminates the 3.6 μm^2^/s state and favors a faster diffusing state at 6.8 μm^2^/s.

### Formation of the oligomeric SctQ complex is dependent on expression of SctQC

Previous work in *Yersinia* has shown that elimination of the internal translation initiation site through a mutation in the *sctQ* coding sequence that replaces the methionine residue at position 218 with an alanine (M218A) results in a secretion-deficient phenotype and that wild-type secretion levels can be restored upon expression of SctQc *in trans* (18, 26). We therefore hypothesized that the *D* ~ 4 μm^2^/s diffusive state measured for eYFP-SctQ in wild-type cells and pYV^−^ background is due to a molecular complex containing SctQ and its C-terminal fragment SctQc (26, 27, 46). To test this hypothesis, we utilized the *eyfp-sctQM218A* coding sequence to express full length eYFP-SctQ, but not SctQC, in the *Y. enterocolitica* pYV^−^ strain (18).

In the absence of SctQC, the distribution of apparent diffusion coefficients of eYFP-SctQ_M218A_ was best fit with four diffusive states (**Fig. 4B, Table 1**). The previously observed diffusive state (D ~ 4 μm^2^/s) of eYFP-SctQ is absent upon elimination of SctQC expression. We instead observe a different, faster moving diffusive state (D = 6.8 μm^2^/s), which is not observed for eYFP-SctQ in either wild-type or pYV^−^ genetic backgrounds. We therefore assign the *D* = 6.8 μm^2^/s diffusive state to a homo-oligomeric SctQ species. Indeed, an oligomeric SctQ-only species was previously detected by co-immunoprecipitation in the absence of SctQc (27). We conclude that the presence of SctQc enables the formation of a well-defined oligomeric SctQ:SctQc complex, which diffuses at *D* ~ 4 μm^2^/s and forms spontaneously in *Y. enterocolitica* in the absence of other T3SS proteins.

The existence of an oligomeric SctQ:SctQC complex is supported by several reports in the recent literature. Bzymek *et al.* expressed *Y. pseudotuberculosis* SctQ and SctQC in *E. coli* and co-purified a SctQ:SctQC complex with 1:2 stoichiometry (*MW* = 52.8 kDa) (26). McDowell *et al.* found that *S. flexneri* SctQ:SctQC was further able to form higher order oligomers consisting of up to six copies of the minimal 1:2 complex and that formation of such higher order oligomers was essential for secretion (46). The quaternary structure of *in situ* injectisomes, recently provided by cryo-electron tomography of *Shigella flexneri* and *Salmonella enterica* minicells (10, 21, 22), allows us to further speculate on the stoichiometry of the oligomeric SctQ:SctQC complex. The sub-tomogram averaged injectisomes displayed six cytoplasmic “pods” that, on one side, attach to the membrane-embedded ring of the needle complex and, on the other side, to hexameric SctN. The observed protein densities of the pods are large enough to contain a tetramer of SctQ proteins, such that 24 SctQ molecules (six tetramers) can be bound to the injectisome at any one time. This value agrees with previously estimated numbers of fluorescently-labelled SctQ proteins within a single diffraction-limited fluorescent focus/cluster (N = 22 ± 8, 28 ± 7, ~24 ± 5) (18, 19, 45). Based on the available data to date, we speculate that the *D* ~ 4 μm^2^/s diffusive state observed in wild-type *Y. enterocolitica* cells is the oligomeric SctQ:SctQC complex that consists of four SctQ and eight SctQC subunits.

### SctQ and SctL co-diffuse as a complex in the cytoplasm

Our single-molecule tracking measurements of eYFP-SctQ in wild-type *Y. enterocolitica* reveal the presence of a slowly diffusing *D* ~ 1 μm^2^/s species with a population fraction of 17%. The increased abundance of this diffusive state compared to pYV^−^ cells could indicate the regulated formation of a slowly diffusing high-molecular weight complex. Indeed, high molecular weight complexes with estimated molecular weights ~1 MDa containing SctQ, SctL, SctK, and SctN have been previously identified in pull-downs and size-exclusion chromatography (11, 16). Given the established interactions between SctK-Q-L-N (12–15), we hypothesized that the *D* ~ 1 μm^2^/s diffusive state consists, at least partially, of a complex containing not just SctQ:SctQC, but also SctK, SctL, and possibly SctN (see **Fig. 1**).

To test whether the *D* ~ 1 μm^2^/s or *D* ~ 4 μm^2^/s diffusive states of eYFP-SctQ are complexes that contain SctL, we used allelic-replacement of the native SctL coding sequence on the pYV virulence plasmid to express the PAmCherry1-SctL fusion protein in wild-type *Y. enterocolitica*. Fitting the apparent diffusion coefficient distribution of PAmCherry1-SctL revealed a substantial stationary population (D < 0.50 μm^2^/s) with a population fraction of 56%. Stationary PAmCherry1-SctL trajectories localized preferentially near the cell boundary (**Fig. S7F**). Based on the same arguments made for eYFP-SctQ, we assign the stationary PAmCherry1-SctL trajectories to injectisome-bound molecules. Fitting the non-stationary components of the apparent diffusion coefficient distribution revealed the presence of five diffusive states (**Fig. 5A, Table 1**). The only observed diffusive states in common between eYFP-SctQ and PAmCherry1-SctL in wild-type *Y. enterocolitica* is the *D* ~ 1 μm^2^/s state. The presence of this common state raises the possibility that these two proteins co-diffuse as a high-molecular weight complex in the cytosol of living cells.

**Fig. 5.**
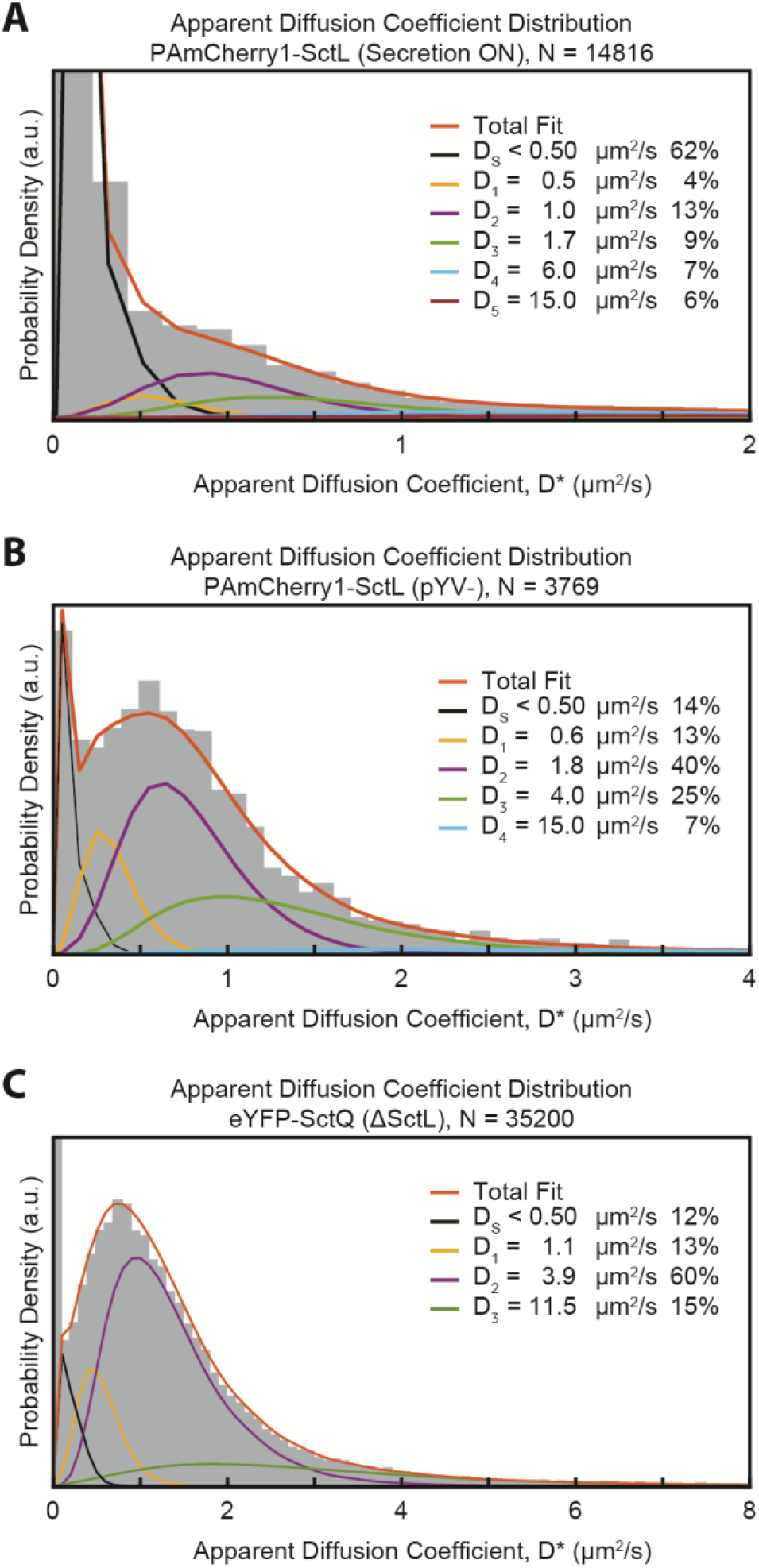
Comparison of PAmCherry1-SctL in the wild-type vs the pYV^−^ strain. (A) Fitting of the apparent diffusion coefficient distribution for PAmCherry1-SctL shows the majority of the population in a stationary state. (B) In the absence of other T3SS proteins the stationary state is depleted, and PAmCherry1-SctL favors faster diffusive states. (C) Apparent diffusion coefficient fitting of eYFP-SctL in a ΔSctL background. Note that the peak near D*=0 is not fit well in this case due to large bin-sizes used in the histogram. We therefore assess the quality of fit using the cumulative distribution function shown in **Fig. S3E** (see also Data Fitting in Materials and Methods).

To test whether the *D* ~ 1 μm^2^/s diffusive state of PAmCherry1-SctL is dependent on the presence of SctQ and other T3SS proteins, we expressed PAmCherry1-SctL in the pYV^−^ strain. In this genetic background, the apparent diffusion coefficient distribution is fit by four diffusive states (**Fig. 5B, Table 1**). A majority of the stationary component is lost in the absence of other T3SS proteins and the population fractions are redistributed to other cytosolic states. Notably absent is a diffusive state near 1 μm^2^/s, indicating that the presence of other T3SS proteins is required for the formation of this state.

As an additional control, we analyzed the diffusive states of eYFP-SctQ in a *ΔsctL* background (**Figure 5C**). The obtained diffusive states, closely resemble those found in the pYV-background, namely, a predominant population in a *D* ~ 4 μm^2^/s diffusive state (60%), in addition to two smaller populations, one in a *D* ~ 12 μm^2^/s diffusive state (15%), and one in a *D* ~ 1 μm^2^/s diffusive state (13%). We assign the fast diffusive state to monomeric eYFP-SctQ, as before. The molecular species responsible for the limited amounts of the *D* ~ 1 μm^2^/s diffusive state remains unclear, but the presence of this state is consistent with the results obtained for eYFP-SctQ and eYFP-SctQM218A in the pYV-backgrounds. The higher population fraction of eYFP-SctQ in wild type cells could thus be due to two different high molecular weight complexes that diffuse at the same rates. The slightly increased population fraction of the stationary component compared to those in the pYV-background could indicate limited binding of eYFP-SctQ to membrane-embedded injectisome precursors, which are known to assemble in absence of SctL(21).

The existence of a high molecular weight complex containing SctQ, SctQC, SctK, SctL, and SctN is consistent with previous FCS measurements showing that the population-averaged diffusion rate of eGFP-SctQ and eGFP-SctL increased when SctN was deleted (19). It is therefore possible that the *D* ~ 1 μm^2^/s diffusive state observed in our work is due to the cytosolic presence of a large supramolecular complex that contains six SctK-Q-L pods, which are each connected to a central hexameric ATPase. Such a supramolecular complex would have a molecular weight of ~2 MDa, assuming previously estimated stoichiometries (18, 19, 21, 22, 45), and could form either through the stepwise assembly in the cytoplasm or upon concerted dissociation from membrane-embedded injectisomes. An alternative explanation is that the *D* ~ 1 μm^2^/s diffusive state represents a single pod that is connected to SctL and possibly SctK and SctN. Even though the Stokes-Einstein relation is not expected to hold in the crowded cytoplasm of living bacterial cells (47, 48), it is not clear how the rather modest addition of SctL (2×25 kDa), SctK (24 kDa), and SctN (48 kDa) subunits to an existing SctQ4:SctQC8 complex (330 or 220 kDa with and without the eYFP tag, respectively) could change its diffusion coefficient by a factor of four. Future work to determine the complete molecular composition of the species responsible for the *D* ~ 1 μm^2^/s diffusive state will help address this question.

### Induction of T3SS secretion alters diffusion behaviors of select cytosolic species

FRAP measurements established that, under secreting conditions, injectisome-bound eGFP-SctQ was dynamically replaced by new cytosolic proteins within t½ ~ 70 seconds (t½ ~ 135 seconds in nonsecreting *Y. enterocolitica* cells) (18). Additionally, FCS measurements found that the population-averaged diffusion rates of SctK, SctQ, and SctL correlate with the secretion state (19). Our results show that the T3SS proteins SctQ and SctL form high molecular weight complexes in the cytosol and that formation of these states is dependent on the presence of other T3SS proteins. Together, these findings suggest the possibility that the type 3 protein secretion pathway is, at least in part, regulated from within the cytosol. We therefore asked whether the diffusion rates or population fractions of SctQ and SctL diffusive states correlate with the secretion state of the cells.

To compare the diffusion behaviors of eYFP-SctQ and PAmCherry1-SctL in secretion ON vs. OFF states we used the fact that the *Y. enterocolitica* T3SS can be switched between secretion ON and OFF states by the addition of EDTA or CaCl_2_, respectively (9). We observed that stimulation of secretion increased the mean apparent diffusion coefficient for mobile molecules (*D\** > 0.15 μm^2^/s) from 1.43 μm^2^/s to 1.69 μm^2^/s for eYFP-SctQ and from 0.86 μm^2^/s to 1. 10 μm^2^/s for PAmCherry1-SctL, which is consistent with recent FCS and earlier 2D-PALM measurements (18, 19). (**Table S3**).

The apparent diffusion coefficient distribution of eYFP-SctQ in secretion OFF conditions was fit with three diffusive states (**Fig. 6A, Table 1**), while that of PAmCherry1-SctL was fit with four diffusive states (**Fig. 6B, Table 1**). The diffusion coefficients obtained for eYFP-SctQ are similar between secretion ON and OFF conditions, suggesting that the oligomerization states of SctQ are not altered upon induction of secretion. However, the relative population fractions do change, suggesting that the relative abundances of SctQ containing complexes are regulated. For PAmCherry1-SctL on the other hand, there seems to be a more complex rearrangement among diffusive states of low abundance. Nonetheless, the presence of the D ~ 1 μm^2^/s diffusive state for both eYFP-SctQ and PAmCherry1-SctL under secretion ON and secretion OFF conditions indicates that the cytosolic interaction between SctQ and SctL is robustly present in both secreting and non-secreting cells. Furthermore, we observe a similar decrease in the relative population fractions of the *D* ~ 1 μm^2^/s diffusive states upon induction of secretion, namely a 26% decrease for eYFP-SctQ and a 24% decrease for PAmCherry1-SctL. These results support our conclusion that SctQ and SctL co-diffuse as high molecular weight complex in the cytosol and suggest that the strength of the SctQ and SctL interaction is not subject to functional regulation of the T3SS ON/OFF switch.

**Fig. 6.**
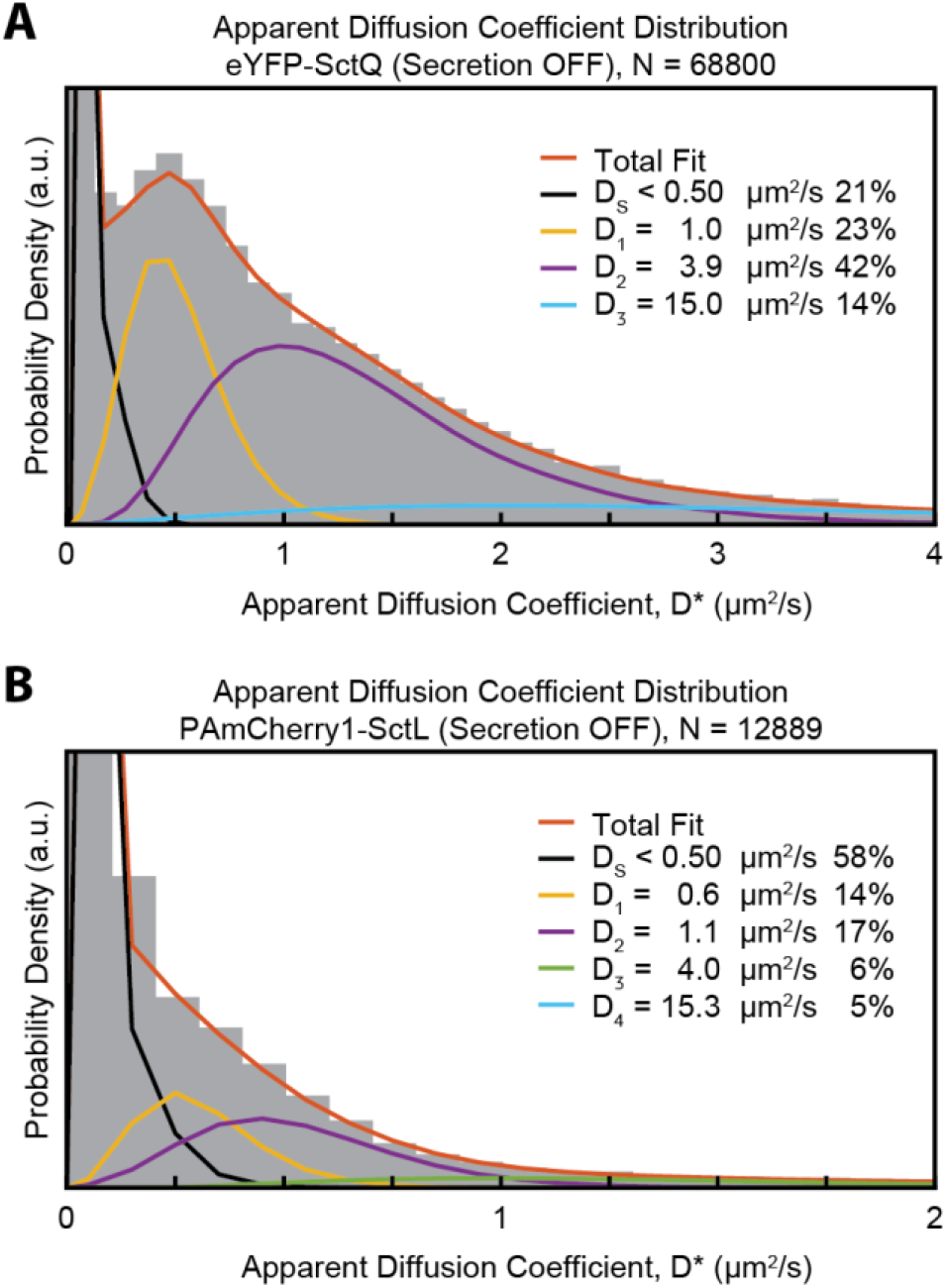
Apparent diffusion coefficient distribution fits for eYFP-SctQ and PAmCherry1-SctL under secretion OFF conditions. (A) Fitting of secretion OFF conditions for eYFP-SctQ. While the individual diffusion coefficients remain the same as for secretion ON conditions, there is an overall population shift towards slower diffusing states. (B) Fitting of secretion OFF conditions for PAmCherry1-SctL. As is the case for eYFP-SctQ, there is a shift towards slower diffusing states for secretion OFF conditions.

## Conclusions

While it’s currently unknown how the cytosolic sorting platform proteins exert their role in the function of the T3SS, it has become clear that their ability to form dynamic cytosolic complexes is linked to function. However, these complexes have remained poorly defined. Using high-throughput 3D single-molecule tracking in living bacterial cells, we resolved the diffusive states of eYFP-SctQ and PAmCherry1-SctL and quantified their respective population fractions. This allowed us to analyze the cytosolic complex formation of the two central sorting platform proteins SctQ and SctL. Our data are consistent with a model in which cytosolic SctQ undergoes dynamic assembly and disassembly steps to interconvert between at least three distinct molecular species that diffuse at different rates. SctQ monomers diffuse freely in the cytosol (D ~ 14 μm^2^/s) or self-assemble into oligomeric SctQ:SctQC complexes (D ~ 4 μm^2^/s) with the help of the C-terminal fragment SctQC. The self-assembly process does not require the presence of other T3SS proteins as evidenced by the observation that a *D* ~ 4 μm^2^/s diffusive state is present in both wild-type and pYV^−^ backgrounds. The SctQ:SctQC complexes do not yet contain SctL, because SctL molecules tracked in wild-type cells do not exhibit a *D* ~ 4 μm^2^/s diffusive state. However, it remains a possibility that SctQ:SctQC complexes in wild-type cells contain SctK, which has been shown to interact with SctQ (12–15). Ultimately, the SctQ:SctQC complexes further associate with SctL, SctK, and possibly SctN to form hetero-oligomeric high molecular weight complexes that diffuses at a rate of *D* ~ 1 μm^2^/s. The presence of a *D* ~ 1 μm^2^/s diffusive state in both the eYFP-SctQ and PAmCherry1-SctL data suggest that SctQ and SctL co-diffuse as part of high molecular weight complexes, but it remains to be determined whether these complexes are individual sorting platform pods or large supramolecular complexes containing six pods that are each connected to a central hexameric ATPase.

At this stage, it is still unclear which of the defined complexes detected in this manuscript plays which role in the secretion process. However, our work reveals two distinct fundamental changes in the architecture of the cytosolic complexes upon activation of the T3SS: either the relative abundances of diffusive states (for SctQ) or the diffusive states themselves (for SctL) are altered upon induction of secretion suggesting a delocalized mechanism of T3SS functional regulation. The diffusive-state-resolved insights add to a growing body of evidence that points to the existence of a dynamic network of cytosolic interactions among structural injectisome proteins and complexes thereof. The activity and substrate selectivity of T3SSs may thus not be programmed into the quaternary structure of the injectisome itself, but instead established in the cytosol through dynamic interactions between T3SS components. Given the *in situ* morphology of injectisomes, it is tempting to speculate that molecular turnover at the injectisome is the result of dynamic binding and unbinding events of individual sorting platform pods and/or entire injectisome cytoplasmic complexes. Such dynamics may help regulate the secretion activity of the T3SS or enable the shuttling of secretion substrate:chaperone complexes to the injectisome. Similar mechanisms might be at play to regulate the structurally similar flagellar motor complex. Chaperone:substrate:FliH_2_-FliI complexes (SctL and SctN homologues) have been isolated from cell extracts (49, 50), and FliI, possibly as part of such a complex, exchanges between the cytosol and the flagellar basal body (51). The bound FliI interacts with the switch complex, which is responsible for controlling the direction of the flagellar rotation, through an interaction between FliH and FliN (SctQ homologue) (52, 53). Regulated assembly of cytosolic protein may thus be a widespread mechanism through which the T3SS and similar bacterial secretion systems are functionally regulated. The present work provides a general strategy to resolve and quantify cytosolic complex formation in living bacterial cells.

## Acknowledgments

We thank Linda Columbus and Jim Casanova for critical reading of the manuscript. We also thank Judith P. Armitage for support and stimulating discussions.

## Notes

Supplementary figures can be found online.

J.R., C.R., A.D., and A.G. and designed research; J.R. and A.D. performed research; J.R., C.R., M.Z., C.D., E.C., A.D., and A.G. contributed new reagents/analytic tools; J.R., C.R., and A.G. analyzed data; and J.R., C.R. A.D., and A.G. wrote the paper.

